# A 7-member SNP Assay on the iPlex MassARRAY Platform Provides a Rapid and Affordable Alternative to Typing Major African *Staphylococcus aureus* Types

**DOI:** 10.1101/395079

**Authors:** Justin Nyasinga, Cecilia Kyany’a, Raphael Okoth, Valerie Oundo, Daniel Matano, Simon Wacira, Willie Sang, Susan Musembi, Lillian Musila

**Affiliations:** United States Army Medical Research Directorate - Africa. P.O. Box 621-00606, Nairobi, Kenya; Kenyatta University. P.O. Box 43844-00100, Nairobi, Kenya; Kenya Medical Research Institute, P. O. Box 54840-00200, Nairobi, Kenya; Technical University of Kenya. P. O. Box 52428-00200, Nairobi, Kenya

**Keywords:** *S.aureus*, MRSA, typing, iplex massARRAY, *spa*, MLST, Kenya

## Abstract

**Background:** Data on the clonal distribution of *Staphylococcus aureus* in Africa is scanty, partly due to high costs and long turnaround times imposed by conventional genotyping methods such as *spa* and multilocus sequence typing (MLST) warranting the need for alternative typing approaches. This study applied and evaluated the accuracy, cost and time of using iPlex massARRAY genotyping method on Kenyan staphylococcal isolates.

**Methods:** Fifty four clinical *S. aureus* isolates from three counties were characterized using iPlex massARRAY, *spa* and MLST typing methods. Ten Single Nucleotide Polymorphisms (SNPs) from the *S. aureus* MLST database were assessed by iPlex massARRAY.

**Results:** The iPlex massARRAY assay grouped the isolates into 14 SNP genotypes with 9/10 SNPs interrogated showing high detection rates (average 89%). s*pa* and MLST typing revealed 22 *spa* types and 21 STs that displayed unique regional distribution. *spa* type t355 (ST152) was the dominant type and t2029 and t037 (ST 241) were observed among MRSA strains. MassARRAY showed 83% and 82% accuracy against *spa* and MLST typing respectively in isolate classification. Moreover, massARRAY identified all MRSA strains and a novel *spa* type. MassARRAY had reduced turnaround time (<12 hrs) compared to *spa* (3 days) and MLST (20 days) typing. The iPlex massARRAY cost approximately 18 USD compared to *spa* (30 USD) and MLST (126 USD) typing based on consumable costs/isolate.

**Conclusion:** Upon validation with a larger collection of isolates, iPlex massARRAY could provide a faster, more affordable and fairly accurate method of resolving African *S.aureus* isolates especially in large surveillance studies.

## INTRODUCTION

Antimicrobial resistance in *S. aureus* is a recognized global threat in clinical management of infections caused by the pathogen [1, 2]. The organism, which may exist asymptomatically in healthy individuals, is capable of causing a wide range of clinical syndromes including skin and soft tissue infections [3]. In Kenya, infections caused by *S. aureus* including methicillin resistant strains (MRSA) are widespread [4-6]. Recently, vancomycin resistant strains (VRSA) have been reported in Kenya’s national referral hospital [7].

MRSA and VRSA strains are often associated with multi-drug resistance, the consequences of which include limited therapeutic options, long treatment periods, escalated treatment costs and significantly higher morbidity and mortality rates [2, 7]. Initially MRSA infections were associated with hospital settings such as surgical wards and the use of indwelling devices but community acquired strains (CA-MRSA) surfaced in the 1990s in Australia and North America and have now spread to all parts of the world [2, 8]. Other MRSA lineages (LA-MRSA) have been associated with livestock infections.

To curb drug resistance, adequate surveillance data regarding the emergence, dissemination and distribution of drug resistant strains is required. *spa* and MLST typing have become the most common molecular methods for characterizing the epidemiology of MRSA strains in surveillance studies [9, 10]. s*pa* typing is a sequence based technique that identifies the number and types of 24 nucleotide repeats in the variable X region of the staphylococcal protein A (*spa*) gene. The method is moderately discriminatory with interlaboratory repeatability and portability [10]. *spa* typing is faster and less expensive compared to MLST since it is a single-locus based technique. However the method presents challenges in delineating particular STs and cannot be reliably applied in outbreak detection [9]. MLST is a sequence based method that indexes polymorphisms in 400-500bp fragments of seven staphylococcal housekeeping genes. With moderate discriminatory power, interlaboratory portability and reproducibility the method is widely applied in surveillance [9]. However, the method is labour intensive, takes long and is costly with low-throughput given that seven loci are analyzed [9].

Particular strains of *S.aureus* tend to localize in certain geographical regions of the world for example the USA100, USA3OO in North America [2] and EMRSA15 and EMRSA16 in the United Kingdom [11]. In Kenya, only two hospital studies have reported on the molecular epidemiology of *S.aureus.* In one study, Omuse *et al*., reported the presence of major global MRSA clones such as MLST-CC5, 22, 7 and 30 among a heterogeneous collection of *S.aureus* isolates [6]. In the same study, members of MLST-CC5 such as ST1, ST5, ST8, ST241 were predominant. The other study conducted by Aiken *et al*., reported *spa* type t037-ST239 as the dominant strain among MRSA isolates recovered from inpatients in a mid-level hospital [5]. Generally, data on the clonal diversity and distribution of MRSA in Africa is scanty warranting further studies [1, 6, 12].

High costs, long result turnaround times, technical complexity and lack of technical expertise have impeded the adoption of conventional typing methods for surveillance studies especially in resource-limited settings such as Africa [1, 6, 12]. In recent years, SNP genotyping has been proposed as a robust, efficient and low cost approach to epidemiological characterization of *S.aureus* [9, 13]. Robertson *et al*., identified and interrogated by real-time PCR, seven informative SNPs that could resolve Australian MRSA strains into groups that reflected the population structure of the organism [14]. Syrmis *et al*., reported 98% accuracy and reduced assay costs when a multiplexed SNP-based assay on the iplex massARRAY platform was compared to real-time PCR SNP genotyping of a collection of Australian *S.aureus* [13]. Despite the potential of iPlex massARRAY as a low cost, rapid and high throughput method for typing staphylococcal isolates, the technique has not been extensively evaluated against conventional methods like *spa* and MLST typing and against *S. aureus* clones from Africa whose epidemiology has been thought to be unique [12].

In this study, results of a multiplexed iPlex massARRAY assay for a collection of Kenyan *S.aureus* isolates are reported. In addition, comparisons between iPlex massARRAY, *spa* and MLST typing with regard to discriminatory power, assay turnaround times and reagent and consumable costs are presented.

## MATERIALS AND METHODS

### Study population and isolates

This study analyzed 54 archived clinical isolates collected between 2015 and 2016 from a recruited patient population of 221 persons. The isolates were part of an ongoing antimicrobial resistance surveillance project covering hospitals in three geographical regions of Kenya. The 54 isolates included 16 isolates from Nairobi County and 19 isolates each from Kisumu and Kericho Counties. Blood, urine, pus and swabs from skin and soft tissue infections and the throat were sampled from consenting in-and out-patients of all ages visiting the participating hospitals. Standard microbiological techniques were used for culture, isolation and identification of *S.aureus.* Isolate identities were further confirmed by conventional PCR for the *femA* gene. Antimicrobial susceptibility testing was performed using both Siemens MicroScan WalkAway 96 plus® (Siemens Healthcare Diagnostics, Inc., New York, USA) and Kirby Bauer disc diffusion methods. About 87% (47/54) of the isolates were associated with community acquired infections. Skin and soft tissue infections (SSTIs) accounted for 94% (51/54) of the infections while urine specimens represented 5.5% (3/54). Eleven percent (6/54) of the isolates were classified as MRSA. All MRSA isolates were *mecA*-positive and 4/6 had SCC*mec*IV genotype. All the isolates were preserved as glycerol stocks at −8°C.

### Ethical Approval

This study was reviewed and approved by the Kenya Medical Research Institute (KEMRI) Scientific and Ethics Review Unit (KEMRI/SERU/CCR/0061/3448) and by the WRAIR IRB (WRAIR #2089A).

### DNA extraction

All clinical isolates and four reference isolates [ATCC 43300, ATCC CO-34, ATCC 25213 and ATCC 25923 (www.beiresources.org)] were sub-cultured by streaking a loopful of inoculum on Muller Hinton agar plates (Sigma-Aldrich, Missouri, USA) and incubating for 18-24hrs at 37°C. The ZR/Fungal/ Bacterial DNA MiniPrep kit (Zymo Research, California, USA) was used for extraction following manufacturer’s instructions with two modifications: The processing time for the cell disruption step was set at 10 mins and DNA was eluted with 60μl of elution buffer. DNA was quantified using Qubit DsDNA quantification kit (Thermo Fisher Scientific, Massachusetts, USA). For iPlex massARRAY assays, DNA concentrations were normalized to 10ng/μl. All DNA samples were stored at −2°C.

### PCR amplification of the *spa* gene

Published primers were used to amplify the X region of the *spa* gene [15]. Each 25 μl PCR reaction mix included 10.5 μl of sterile nuclease free water, 12.5 μl of Dream Taq mix (Thermo Fisher Scientific, Massachusetts, USA), 0.5 μl (10pmol) of both the forward and reverse primers with 1 μl (approximately 50ng) of template DNA. Cycling was performed in a GeneAmp 9700 PCR System (Applied Biosystems, California, USA) with the following conditions: Initial denaturation at 95°C for 5 mins, and 35 cycles of denaturation at 95°C for 45s, primer annealing at 6°C for 30s, extension at 72°C for 90s followed by a final extension at 72°C for 10 mins. Amplification products were resolved on a 1.5% agarose in 1X Tris Acetate EDTA buffer (Sigma-Aldrich, Missouri, USA) for 60 mins at 95V and visualized with EZ Vision DNA stain (Amresco Inc. Ohio, USA) using a UV-transilluminator (UVP LLC, California, USA).

### PCR amplification of MLST loci

Published primers were used to amplify the MLST loci [16]. Each 25 μl reaction comprised 10.5 μl of sterile nuclease free water, 12.5 μl of Dream Taq mix (Thermo Fisher Scientific, Massachusetts, USA), 0.5 μl (10pmol) of both forward and reverse primers with 1 μl of template DNA (approximately 50ng). Thermocycler conditions were set as: Initial denaturation at 95°C for 3 mins and 35 cycles of strand denaturation at 95°C for 30s, primer annealing at 55°C for 30s and extension at 72°C for 60s and a final extension at 72°C for 10 mins. The amplicons were resolved and visualized as described above for the *spa* gene.

### Sanger sequencing of *spa* and MLST amplicons

*spa* and MLST amplicons were purified using the DNA Clean & Concentrator kit (Zymo Research, California, USA) following manufacturer instructions with a final elution volume of 3° μl. Both forward and reverse strands were sequenced. Each reaction contained 4 μl of sterile distilled water, 2 μl of 5X Big Dye buffer (Applied Biosystems, California, USA), 1 μl of the PCR primers (4 μM), 1 μl of Big Dye terminator mix (Applied Biosystems, California, USA) and 4 μl of amplicon DNA. Cycle sequencing was performed on an ABI 9700 thermocycler with cycling conditions set as: 94°C for 5 mins followed by 3° cycles of 94°C for 15s, 55°C for 30s and 68°C for 2 mins and 30s and a final extension of 68°C for 3 mins. Sequencing fragments were purified using Sephadex G5° resin (Sigma-Aldrich, Missouri, USA) before loading on the Applied Biosystems Genetic Analyzer 3500.

### Iplex massARRAY assays for MLST SNPs

The 10 MLST SNPs used in this study have been published before [13]. Nine of the 10 MLST SNPs assayed were bi-allelic while one (arcC210) was tri-allelic. Amplification and extension primers were designed using Agena Bioscience Assay Design Suite Version 2.0 (Agena Bioscience, Hamburg, Germany). The sequences used to design primers for primary amplification and extension PCR reactions were downloaded from the staphylococcal MLST database, http://www.mlst.net, 20^th^ June, 2016 [16] and are shown in Table 1. All primary PCR primers had a 10-nucleotide tag (**ACGTTGGATG**) on the 5**’** end to exclude them from the 4500-9000 Da mass range of MALDI-TOF MS detection. The primary multiplex PCR reactions were run on a GeneAmp 9700 PCR machine (Applied Biosystems, California, USA). Each reaction mix contained 1x PCR Buffer, 0.1 μM of forward and reverse primers, 4 mM MgCl_2_, 500 μM dNTP mix, 0.5 units of Taq polymerase and 10 ng of DNA template. PCR conditions were set as follows: Initial denaturation at 95°C for 2 minutes, 25 cycles of 95°C for 30s, 56°C for 30s, and 72°C for 60s followed by a final extension at 72°C for 5 mins.

**Table 1:**
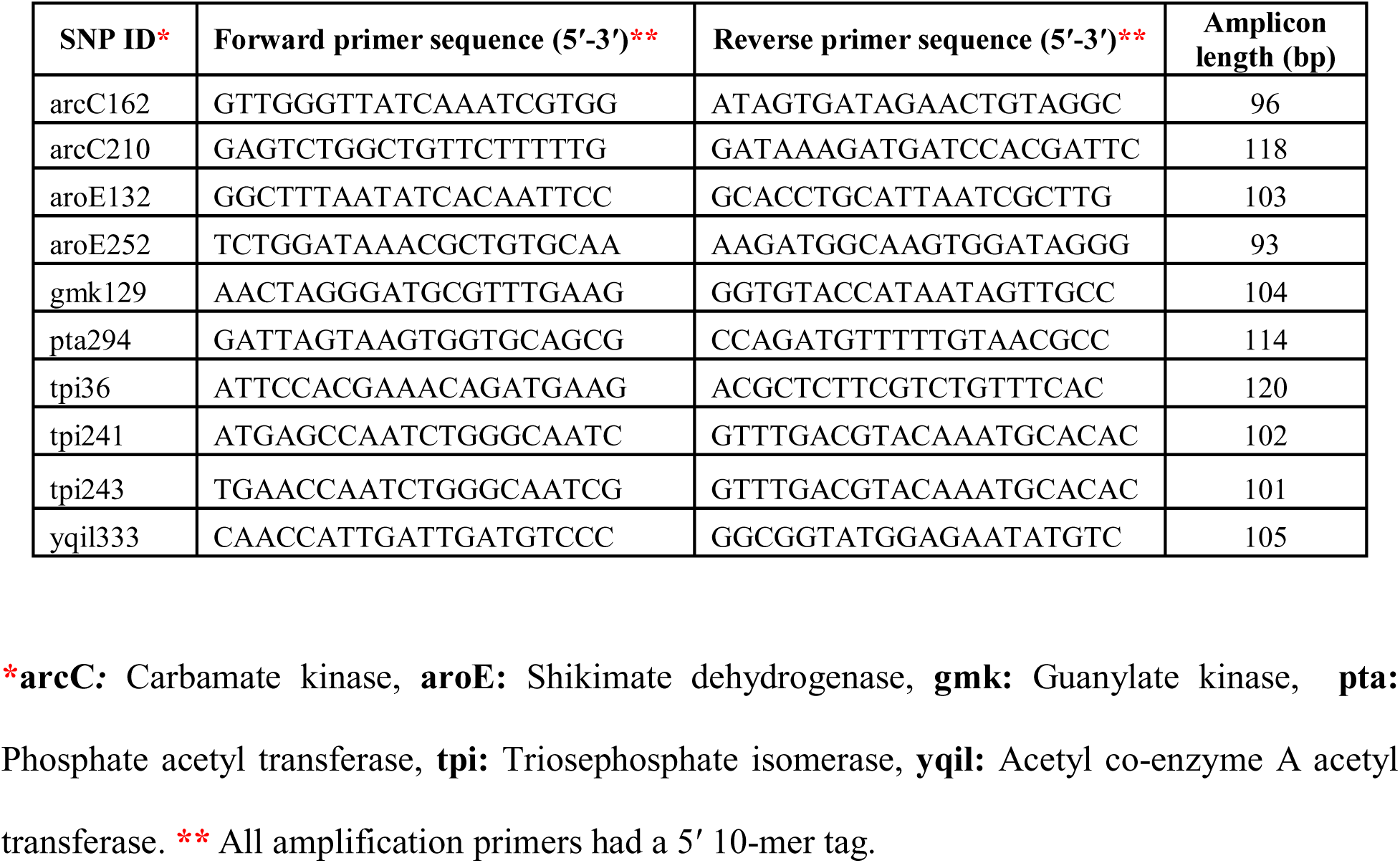
Primary amplification primer sequences for various SNPs and their expected amplicon sizes

Shrimp alkaline phosphatase (SAP) enzyme was used to dephosphorylate unused dNTPs from the primary PCR reaction before the second allele-specific primer extension PCR. For allele specific PCR reactions, 2 μl of the reconstituted extension primer cocktail was added to each reaction well and the following conditions were set: Initial denaturation at 94°C for 30s, 4° cycles of one step at 94°C for 5s with five sub cycles of 52°C for 5s and 8°C for 5s, and a final extension at 72°C for 3 minutes. The sequences and masses for unextended primers (UEPs) and extension products for each SNP and SNP allele are shown in Table 2. Extension products were conditioned using a resin after which 10 nl was dispensed to a 96-well spectroCHIP using the MassARRAY Nanodispenser RS100° (Agena Bioscience, Hamburg, Germany).

**Table 2:**
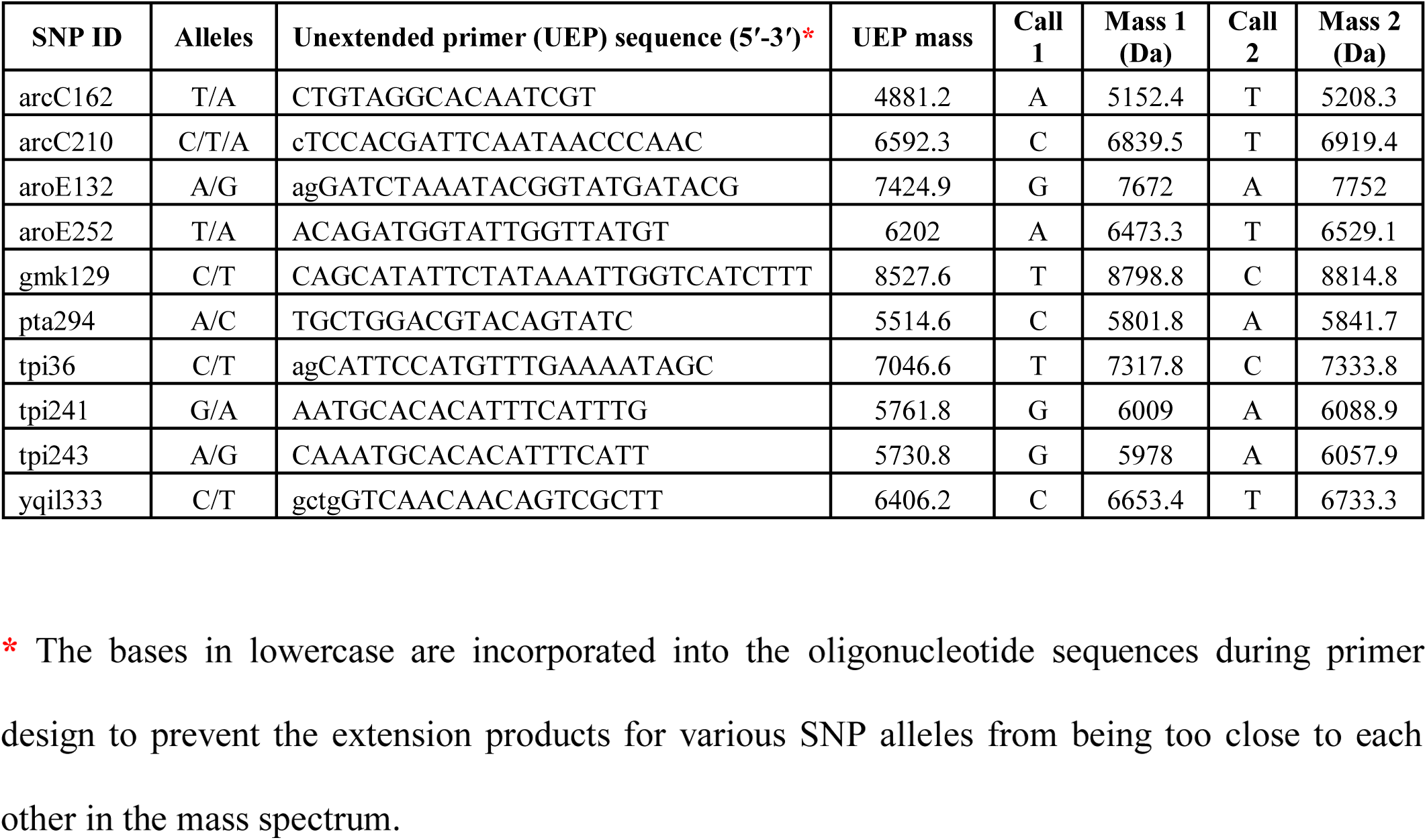
Allele-specific extension primer sequences and expected masses for various SNP alleles

### Costing and time comparisons

Reagent and consumable costs per isolate for *spa* and MLST typing were estimated for various steps such as DNA extraction, primer sequences, PCR amplification, gel electrophoresis, PCR clean up, cycle sequencing, fragment purification and fragment analysis. For iPlex massARRAY, costs for primer sequences, primary and allele-specific PCR amplifications, SAP treatment, sample conditioning and liquid transfer to spectrochips were estimated. Time durations to result generation for the three methods after DNA extraction were also compared.

### Data analysis

Raw sequencing chromatograms were examined using Chromas Version 2.6.2 (Technelysium Pty) before consensus sequences were created using BioEdit Sequence Alignment Editor Version 7.2.5 [17]. *spa* types were assigned using the online s*pa* Type Finder/Identifier Software, *<u>spa</u>*<u>typer.fortinbras.us</u>/ (Fortinbras Research). s*pa* types were further confirmed using the Ridom *spa* Server, *<u>spa</u>*<u>server.ridom.de</u> (Ridom GmbH, Würzburg, Germany) [18]. For MLST loci, consensus sequences were aligned using the online alignment tool, MAFFT Version 7 https://mafft.cbrc.jp/alignment/server/. Sequence types were assigned on the MLST database www.mlst.net [16] using the “Exact or Nearest Match” option. The SpectroAcquire program was used for data acquisition on the MassARRAY Compact Analyzer (Agena Bioscience, Hamburg, Germany) and detection parameters were set at ten laser shots per raster position with a threshold of five good spectra per sample pad.

### Data availability

Raw chromatograms for *spa* and MLST typing as well as data output from the MALDI-TOF MS will be made available upon request.

## RESULTS

### Distribution of *spa* and MLST types

*spa* and MLST typing of the collection of isolates yielded congruent results. *spa* and MLST typing showed 22 *spa* types and 21 STs respectively. The isolates displayed considerable heterogeneity with 16/22 *spa* types and 13/21 sequence types being represented by only one isolate. The three regions showed unique genetic fingerprints with minimal overlaps in isolate composition. Only t355 (ST152) was observed across the three regions with 18 *spa* types and 18 STs associating with specific regions. Kericho region showed greater diversity (12 *spa* types and 10 STs) followed by Kisumu (10 *spa* types and 8 STs) and Nairobi (5 *spa* types and 7 STs). Two novel *spa* types t17841 and t17826 were reported and were both associated with MSSA isolates. Despite the heterogeneity, t355 (ST152), t064 (ST8), t005 (ST22), t2029/t037 (ST241) seemed to dominate. The six MRSA isolates were represented by types: [t2029 (ST241) n=1, t037 (ST241) n=3, t355 (ST152) n=1 and t007 (ST39) n=1].

### Typability and variability of the MLST SNPs

Table 3 summarizes the characteristics of the 10 SNPs analyzed. Nine of the 10 SNPs interrogated were highly typeable with SNP call rates (number of isolates in which a particular SNP was identified as a proportion of the total number of isolates tested) that ranged from 81% - 98.3% (average 89%). One SNP, pta294, was identified in only one isolate and was subsequently excluded from the analyses. Eight of the remaining SNPs were highly variable with allele frequencies (number of isolates positive for a given SNP allele as a proportion of all isolates in which that SNP was identified) ranging from 53% (arcC162) to 84.6% (yqil333).

**Table 3:** Summary of the 10 MLST SNPs analyzed by iPlex massARRAY

| SNP | Polymorphism | SNP call rate <sup>a</sup> | Allele frequency <sup>b</sup> |
| --- | --- | --- | --- |
| arcC162 | T/A | 98.3% (57/58) | **T** 53% (30/57) |
| arcC210 | C/T/A | 86% (50/58) | **T** 60% (30/50) |
| aroE132 | A/G | 91.3% (53/58) | **A** 73.6% (39/53) |
| aroE252 | T/A | 86% (50/58) | **T** 64% (32/50) |
| gmk129 | C/T | 84.5% (49/58) | **C** 67.3% (33/49) |
| tpi36 | C/T | 81% (47/58) | **C** 63.8% (30/47) |
| tpi241 | G/A | 86% (50/58) | **G** 98% (49/50) |
| tpi243 | A/G | 94.8% (55/58) | **G** 60% (33/55) |
| yqil333 | C/T | 89.7% (52/58) | **T** 84.6% (44/52) |
| pta294 | A/C | - * | -* |

\* pta294 was identified in only one isolate and was excluded when generating SNP genotypes.
<sup>a</sup> **SNP call rate:** The proportion of isolates in which a given SNP was identified as a proportion of all isolates tested for that SNP. <sup>b</sup> **Allele frequency:** The number of isolates in which a given SNP allele was identified as a proportion of all isolates in which the SNP was identified.

### Iplex massARRAY SNP genotypes

The combination of nine SNPs for each isolate was used to generate SNP genotypes. Different SNP combinations were created with the aim of finding the smallest number of SNPs with the highest resolution (Data not shown). A 7-member SNP classification that excluded both aroe252 and tpi36 **(arcC162, arcC210, aroE132, gmk129, tpi 241, tpi 243** and **yqil333)** achieved a similar resolution to that of the 9-member SNP profiles. The assay grouped 44/54 isolates into 14 different SNP genotypes. Table 4 shows the regional distribution of various SNP genotypes. In Kericho, nine SNP genotypes were observed among the isolates while Kisumu and Nairobi isolates showed 8 and 4 SNP genotypes respectively. The assay successfully typed 11/16, 16/19 and 17/19 isolates from Nairobi, Kericho and Kisumu respectively while 10 isolates did not have complete SNP profiles. The classifications achieved by iPlex massARRAY reflected the heterogeneity and regional distribution patterns revealed by *spa* and MLST typing.

**Table 4:**
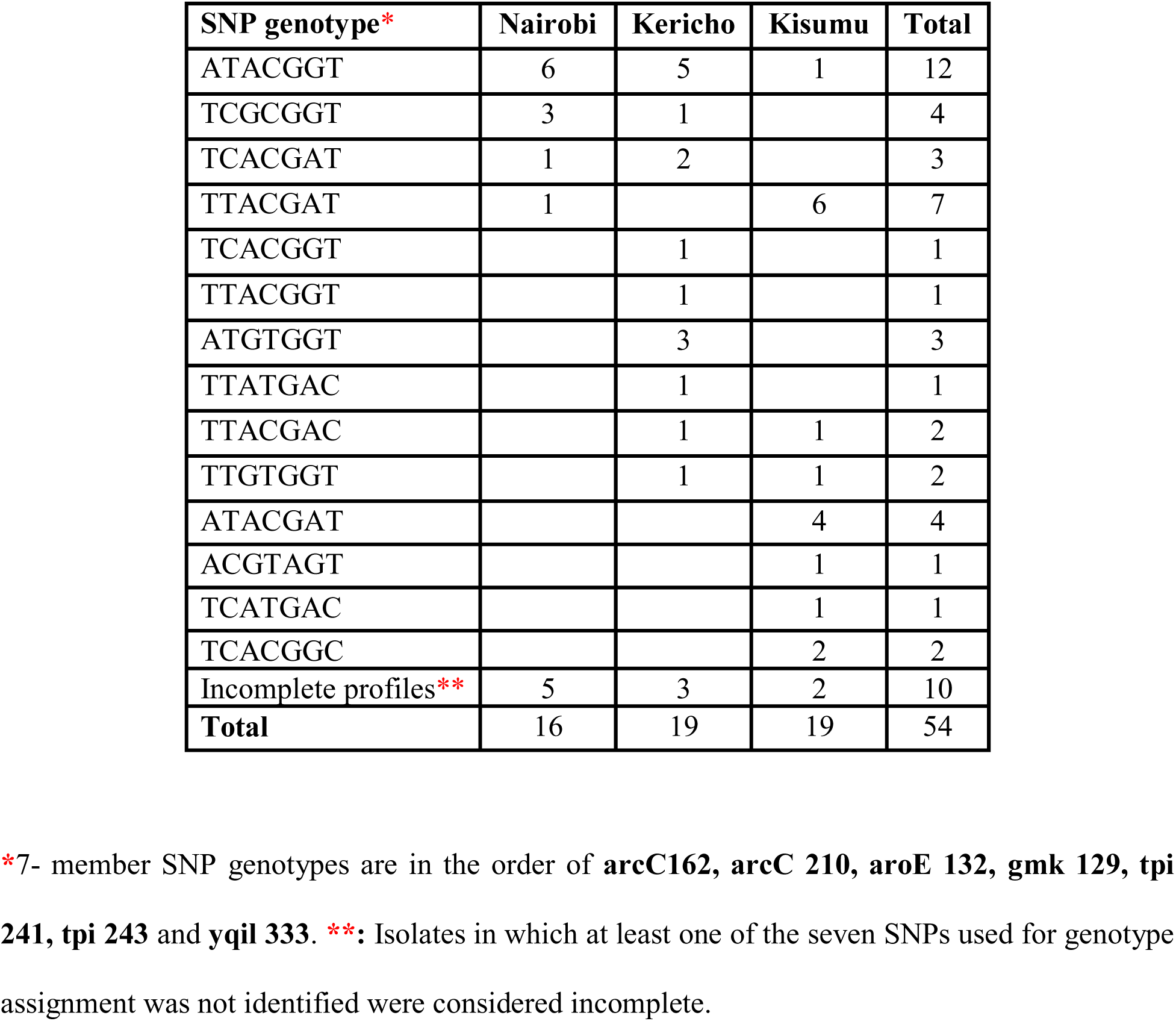
Regional distribution of *S. aureus* isolates by massARRAY SNP genotyping

The 14 SNP genotypes were compared with corresponding *spa* and MLST sequence types for all isolates. Iplex massARRAY showed 83% (34/41) accuracy against *spa* typing and 82% (32/39) against MLST typing in genotype assignment. The SNP genotype **TTACGAC** corresponded to a novel *spa* type t17841 and SNP genotype **TCACGGT** corresponded to another novel *spa* repeat sequence yet to be assigned a *spa* type. iPlex massARRAY identified all MRSA isolates in this collection. One MRSA was of the **ATACGGT** genotype which corresponded to t355. t037 which was the *spa* type of 3 MRSAs and t2029 (1 MRSA) were represented by the **ATACGAT** genotype. **ACGTAGT** corresponded to t007 (one MRSA).

Three discrepancies involving iplex massARRAY were observed: SNP genotype **TCACGGC** could not distinguish between t3772 and t13194 (ST 25 and ST80). SNP genotype **TTGTGGT** could not distinguish between t314/ and t272 (ST121 and 152). Lastly, **TCACGAT** could not distinguish between t084 and t131 (ST 15 and ST1290). Discrepancies involving *spa* types t318/t021 and t037/t2029 were considered minor as such isolates are grouped as members of ST30 and ST241 respectively by MLST (Table 5).

**Table 5:**
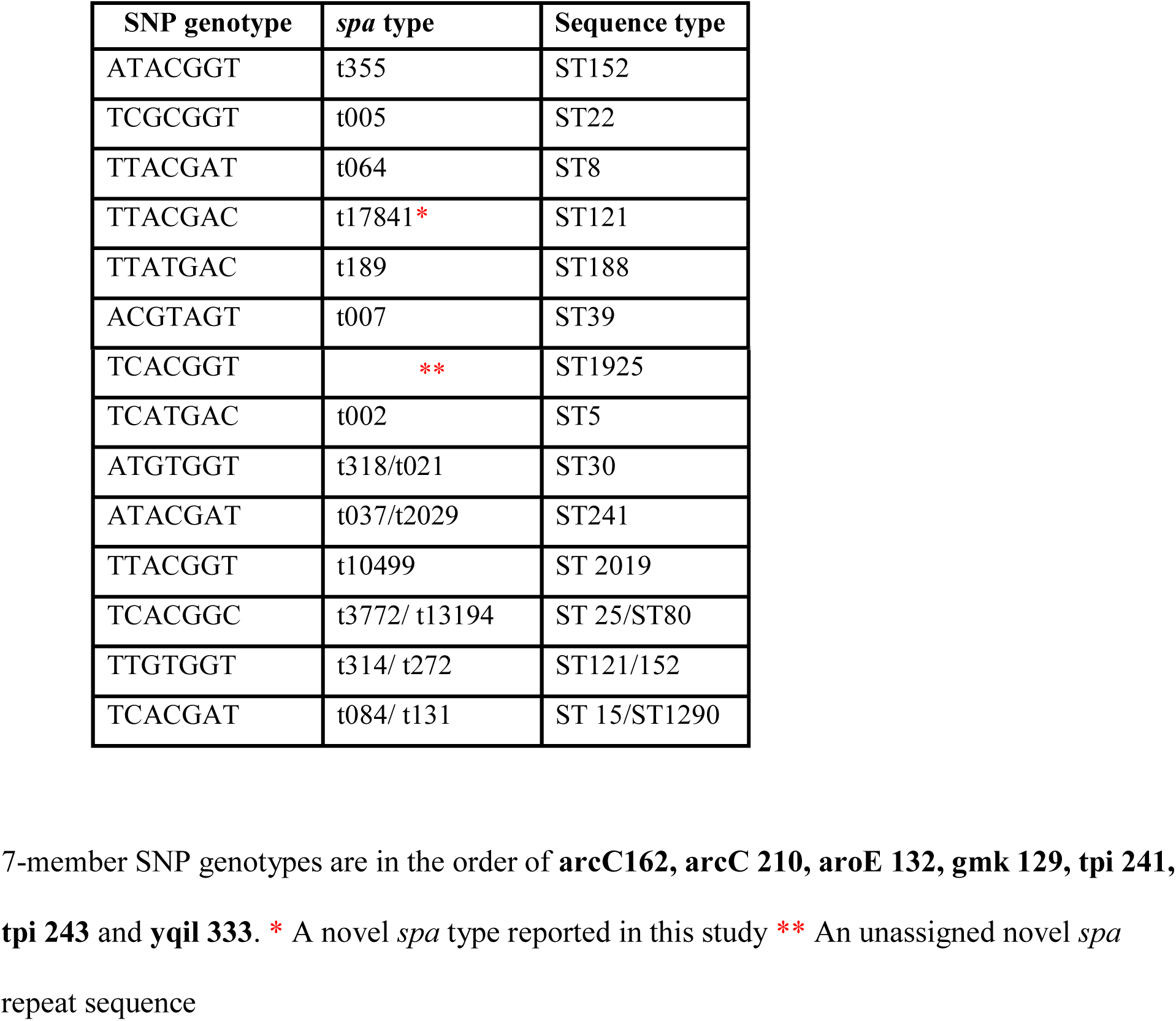
A summary of comparisons between iplex massARRAY, *spa* and MLST typing

### Turnaround time, reagent and consumable costs per isolate

The iPlex massARRAY assay took approximately 1 day compared to *spa* and MLST typing which took approximately 3 and 20 days respectively (Table 6). The reagent and consumable cost of analyzing an isolate for massARRAY was approximately 18 USD compared to *spa* (30 USD/isolate) and MLST (126 USD/isolate) typing for the collection of isolates analyzed (Data not shown).

**Table 6:**
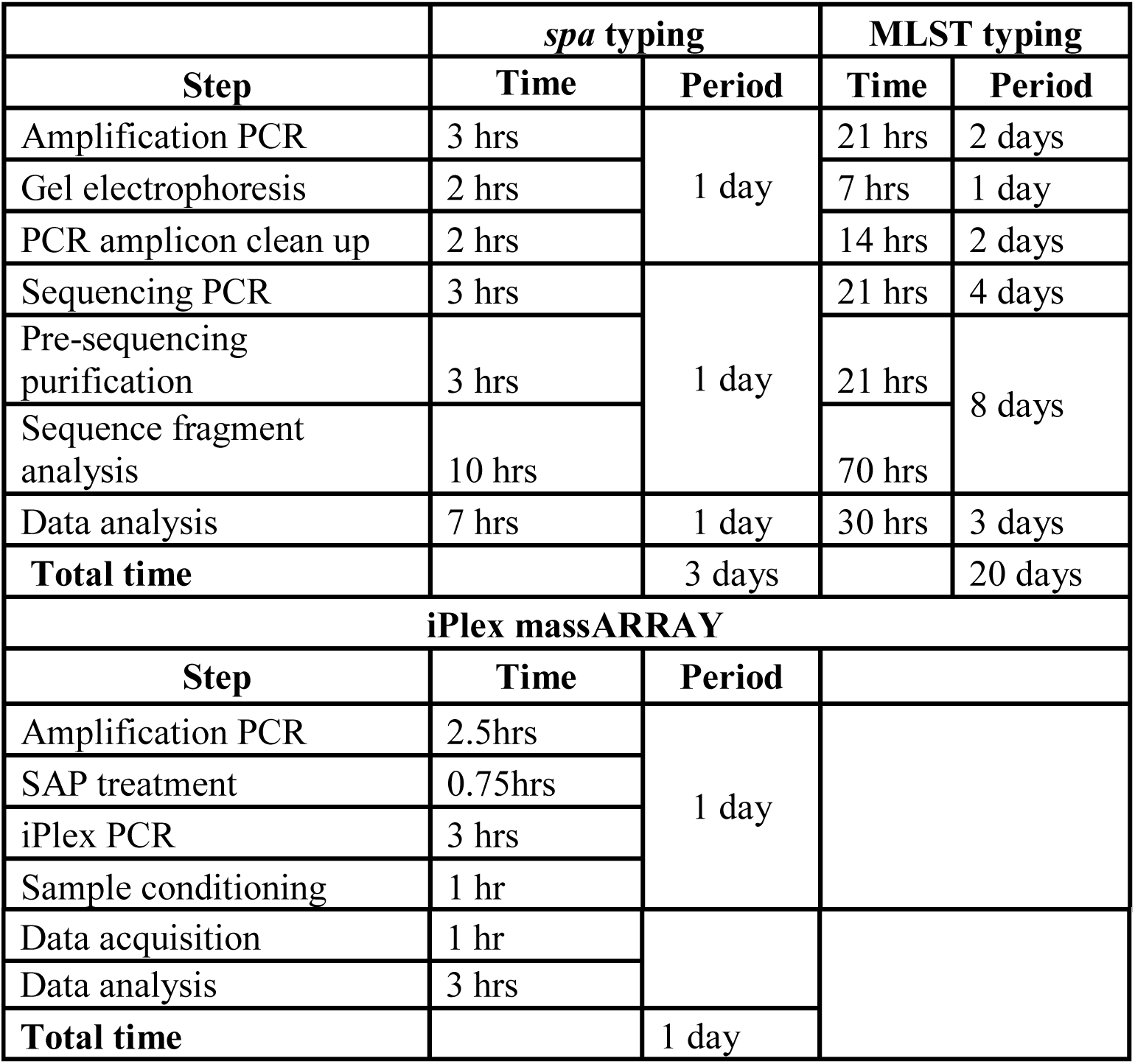
Time-to-result analysis for *spa*, MLST and massARRAY typing methods for the isolates after DNA extraction

|  | *spa* typing |  | MLST typing |  |
| --- | --- | --- | --- | --- |
| Step | Time | Period | Time | Period |
| Amplification PCR | 3 hrs | 1 day | 21 hrs | 2 days |
| Gel electrophoresis | 2 hrs |  | 7 hrs | 1 day |
| PCR amplicon clean up | 2 hrs |  | 14 hrs | 2 days |
| Sequencing PCR | 3 hrs | 1 day | 21 hrs | 4 days |
| Pre-sequencing purification | 3 hrs |  | 21 hrs | 8 days |
| Sequence fragment analysis | 10 hrs |  | 70 hrs |  |
| Data analysis | 7 hrs | 1 day | 30 hrs | 3 days |
| **Total time** |  | 3 days |  | 20 days |
| **iPlex massARRAY** |  |  |  |  |
| Step | Time | Period |  |  |
| Amplification PCR | 2.5hrs | 1 day |  |  |
| SAP treatment | 0.75hrs |  |  |  |
| iPlex PCR | 3 hrs |  |  |  |
| Sample conditioning | 1 hr |  |  |  |

|  |  |  |
| --- | --- | --- |
| Data acquisition | 1 hr |  |
| Data analysis | 3 hrs |  |
| **Total time** |  | 1 day |

## DISCUSSION

Seven MLST SNPs identified 14 unique genotypes and reflected the heterogeneity and distribution depicted by *spa*/MLST typing of a collection of 54 Kenyan isolates. iPlex massARRAY identified all the MRSA isolates used in this study as well as detecting a novel *spa* type. The assay showed comparable resolution to *spa* and MLST typing (83% and 82% respectively). The combination of 7 SNPs achieved similar resolution to that of 9 SNPs initially assayed. Coincidentally, the seven SNPs matched those originally described by Robertson *et al* that could resolve major Australian MRSA strains [14]. A technique that can identify circulating strains within a population using the smallest number of variable loci would offer the benefits of reduced costs and turnaround time. The isolate types identified by massARRAY such as ST152, ST5, ST8, ST15, ST30 and ST241 have been reported in other African countries such as Cameroon, Madagascar, Morocco, Niger, and Senegal [19] and Ghana [20] thereby highlighting the potential of this assay to be universally applied in the African context.

In some instances the assay could not distinguish between two or more *spa*/MLST types, an observation noted by others [14, 21]. The three SNP profiles that could not distinguish certain *spa*/MLST types did not belong to major circulating *spa*/MLST types in the study as 5/6 affected groups were represented by one isolate each, none of which was an MRSA strain. Previously, it has been suggested that increasing the number of SNPs from 7 to 14 can resolve minor *S. aureus* clones [21]. However, 2 of the 10 SNPs (aroE252 and tpi36) assayed did not prove useful in increasing the resolution of the assay while one (pta294 was poorly typeable). While the potential of a larger SNP set to resolve minor clones is recognized, a different set of SNPs in addition to the seven identified here may be required.

MassARRAY demonstrated great capabilities for increased speed and throughput. After optimization, the assay took <12 hours from DNA amplification to generate results for the isolates analyzed in comparison to *spa* (3 days) and MLST typing (20 days). The technique offers greater multiplexing capabilities that can upon effective assay design be used to detect other genes to answer clinically important questions about a pathogen such as virulence and resistance [13] a feature that is absent both in *spa* and MLST typing methods.

MassARRAY showed considerable reduction in assay costs in comparison to *spa* and MLST typing. Estimated reagent and consumable cost/isolate was 18USD compared to *spa* and MLST typing (30USD and 126 USD respectively). In one study, when the iplex massARRAY assay costs were compared to SYBR green real time PCR, there was a 60% reduction in reagent costs [13]. Trembizki *et al*. noted an approximately 30% reduction in cost compared to performing a full MLST analysis [22]. The fairly high initial costs for the massARRAY system are justified by the short turnaround times, multiplexing, automation and throughput capabilities which can support multiple large scale studies concurrently thereby contributing to massive amounts of data.

As the massARRAY technology is increasingly being adopted, studies utilizing its application in bacterial genotyping are being reported. Trembizki *et al* developed, applied and validated a 14-member SNP assay on the massARRAY platform for genetic characterization of *Neisseria gonorrhea* isolates in Australia [22]. Subsequently, the assay was used for a large scale AMR surveillance study of *Neisseria gonorrhea* [23].

The assay is promising particularly in long-term surveillance studies where it is impractical to perform *spa* and MLST typing on the hundreds to thousands of isolates recovered from such studies. An ideal approach would be to perform an iPlex massARRAY analysis as a first-step molecular screen to identify isolates with unusual SNP genotypes or those with which discrepant results are recognized. A full *spa* or ST determination could then follow to definitively resolve such isolates.

There were instances for example with pta294 where particular SNPs could not be called. A possible explanation for this could be sequence variation in the primer targets for the primary PCR reactions, an observation that has been noted elsewhere [13, 22]. The quality of DNA can potentially affect the success of the assay [22]. However, the DNA used for these experiments was of high quality as extraction was done using a commercial kit and DNA concentrations were measured and normalized.

The lack of a database for MLST SNPs that can be used for matching particular SNP profiles to known STs or *spa* types [22] is a major limitation. This is partly due to the fact that assays on this platform have not been rigorously evaluated against known STs/*spa* types from an international collection of isolates. However, with increased validation and adoption, it should be possible to have an online MLST-style platform where SNPs can be submitted for inter-laboratory comparability of data.

Compelling epidemiological conclusions cannot be drawn as the study analyzed a modest collection of 54 isolates, a small proportion of which constituted MRSA isolates. A larger collection of isolates from diverse regions and clinical syndromes would give not only a reliable reflection of the epidemiology of MRSA in Kenya but also serve to highlight the utility of iplex massARRAY for surveillance.

In conclusion, seven SNPs derived from the MLST loci provided comparable discriminatory power for resolving a heterogeneous and regionally unique collection of Kenyan clinical *S.aureus* isolates including MRSA strains. The iplex massARRAY demonstrated advantages of reduced turnaround time and assay costs in relation to two conventional typing methods. With increased validation, the assay should serve as a complement to existing typing methods especially in staphylococcal AMR surveillance studies.

## Acknowledgements

This work was funded by the Armed Forces Health Surveillance Branch (AFHSB) and its Global Emerging Infections Surveillance (GEIS) Section. PROMIS ID: 20160270153 FY17. JN received a student research grant from the Kenya National Research Fund. We acknowledge the use of the *S. aureus* MLST database which is located at Imperial College London and is funded by the Wellcome Trust. This publication made use of the *spa* typing website (http://www.spaserver.ridom.de/) that is developed by Ridom GmbH and curated by SeqNet.org (http://www.SeqNet.org/). We appreciate technical support from Luiser Ingasia and the MDR Laboratory. We thank the study participants for the AMR Surveillance Project. This article has been published with permission from the Director, KEMRI.

## Disclaimer

Material has been reviewed by the Walter Reed Army Institute of Research. There is no objection to its presentation and/or publication. The opinions or assertions contained herein are the private views of the author, and are not to be construed as official, or as reflecting true views of the Department of the Army or the Department of Defense. The investigators have adhered to the policies for protection of human subjects as prescribed in AR 70-25.

## Author contributions

**JN:** Study design, strain typing, data analysis and manuscript write up and review; **LM:** Study design, data analysis, manuscript write up, review and proof reading; **CK:** DNA extraction, strain typing and manuscript review; **RO:** MassARRAY assay design and execution, data analysis and manuscript review. **VO, DM** and **SW** performed bacterial culture and identification, AST testing and manuscript review; **WS** and **SM** offered technical consultation and manuscript review.

